# Top-down modulation of visual cortical stimulus encoding and gamma independent of firing rates

**DOI:** 10.1101/2024.04.11.589006

**Authors:** Christopher M. Lewis, Thomas Wunderle, Pascal Fries

## Abstract

Neurons in primary visual cortex integrate sensory input with signals reflecting the animal’s internal state to support flexible behavior. Internal variables, such as expectation, attention, or current goals, are imposed in a top-down manner via extensive feedback projections from higher-order areas. We optogenetically activated a high-order visual area, area 21a, in the lightly anesthetized cat (OptoTD), while recording from neuronal populations in V1. OptoTD induced strong, up to several fold, changes in gamma-band synchronization together with much smaller changes in firing rate, and the two effects showed no correlation. OptoTD effects showed specificity for the features of the simultaneously presented visual stimuli. OptoTD-induced changes in gamma synchronization, but not firing rates, were predictive of simultaneous changes in the amount of encoded stimulus information. Our findings suggest that one important role of top-down signals is to modulate synchronization and the information encoded by populations of sensory neurons.

## Introduction

The brain integrates external signals arising from the sensory periphery with internal signals related to attention, expectation, motor actions, previous experience, and the subject’s current goals^1,2^. The integration of external ‘bottom-up’ and internal ‘top-down’ information enables flexible, state-dependent processing of sensory stimuli and forms the basis of adaptive behavior^3–5^.

Neuroscience has classically characterized the response properties of sensory neurons in relation to the features of external stimuli due to their experimental accessibility^6–9^. While this procedure has yielded much information about the bottom-up processing of sensory signals, the advent of cognitive neuroscience and the introduction of well-controlled psychophysical tasks, first in humans^10^ and non-human primates^11,12^ and later in rodents^13^, fish^14^, and insects^15^ has increased our awareness for the state-dependent, top-down modulation of neurons, even in primary sensory areas. The operationalization of cognitive processes in psychophysical tasks designed to isolate distinct top-down influences has yielded an appreciation for the sensitivity of sensory neurons in the thalamus and cortex to top-down modulatory influences^16–21^. However, operationalizing top-down influences in this manner likely engages many distinct processes simultaneously in the awake, behaving animal, and some may be impossible to disentangle because psychological factors do not cleanly distinguish the underlying neuroanatomy conveying top-down signals.

Top-down influences arise from many distinct sources, including higher-order sensory areas, association and motor cortices, as well as higher-order thalamic and neuromodulatory nuclei^22–27^. These diverse sources might have distinct effects on their target population, and experimental perturbations of specific neurons, nuclei, or brain areas can help elucidate their contributions to the top-down modulation of sensory coding.

Within the visual cortex, the feedforward and feedback projections that convey bottom-up and top-down influences are largely anatomically segregated in terms of their laminar origins, terminations, as well as in the characteristics of their synapses^28–31^, and these differences can be used to define a hierarchy of visual areas^32,33^. While the visual hierarchy of the cat and primate are the paradigmatic examples of hierarchical cortical processing^28,32,33^, hierarchical processing in sensory systems appears conserved across mammals, with mice exhibiting hallmarks of a visual hierarchy^34–37^. Thus, one prominent source of top-down information in sensory processing are the extensive anatomical feedback projections from higher-order to early sensory areas^32,38^. These extensive feedback projections within the visual hierarchy may, in addition to attention and expectation signals, convey visual spatial or temporal context, or predictions that enrich the information available in the classical receptive field of primary visual cortex neurons^38–41^.

We sought to investigate the effects of top-down signals from higher visual cortex on primary visual cortex in the lightly anesthetized cat. The cat visual system is highly developed, exhibiting many of the hallmark features of the primate visual system, such as a well-defined hierarchy of visual areas with tight functional and anatomical homologies to the primate^33,42^ and has long served as a model system for sensory processing^8^ and the laminar organization of sensory systems^28,43^. As in the primate, cat V1 neurons receive extensive feedback connections from higher-order sensory and association areas, which can modulate their response properties^44,45^.

We chose to target a higher-order area in the ventral stream of the cat, area 21a, a homologue to area V4 in the primate^42^ (Fig. 1A). Area 17 and 21a have bidirectional monosynaptic connections that show high spatial modularity and specificity^46^ and that are prototypical in terms of the laminar distribution^47,48^ of the sources and recipients of their feedforward and feedback connections^33,49^ (Fig. 1B). Previous studies have investigated the functional effects of silencing area 21a on the sensory responses of area 17 neurons by cooling area 21a^44,45^, finding reduced tuning and spatial integration. To date, these investigations have focused on top-down effects on the firing rate of V1 neurons. However, top-down signals might have diverse effects on the response properties of neurons and neuronal populations in sensory cortices, potentially affecting not only the firing rates or tuning of single neurons, but also the synchronization of neurons and populations and the information they encode. In awake primates, top-down signals of expectation and attention have been shown to modulate both gamma-band synchronization and firing rates in primary visual cortex^21,50–55^. Notably, behaviorally isolated top-down signals have been shown to both increase and decrease gamma-band synchronization and firing rates in the primary visual cortex^21,50,53^.

**Figure 1:**
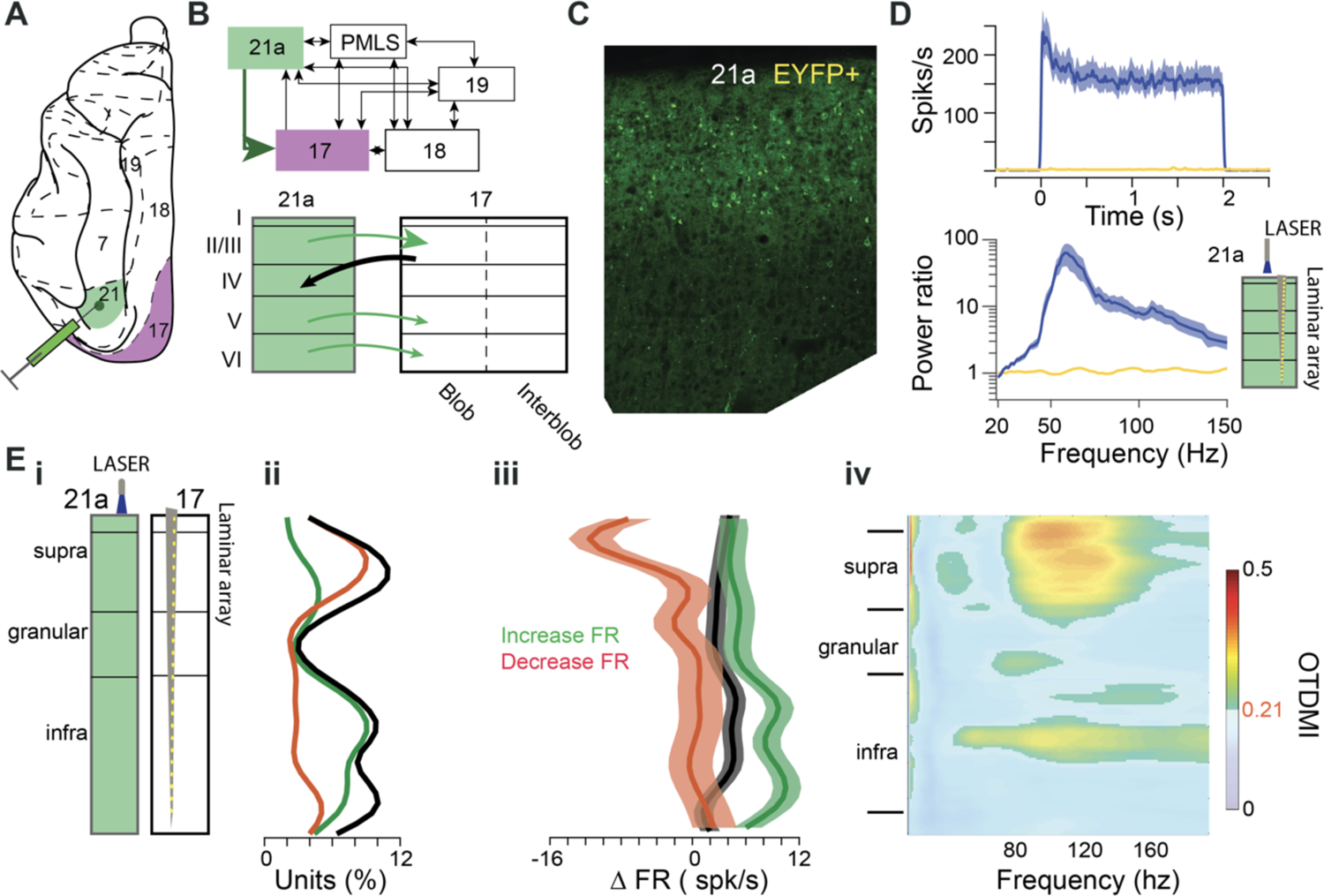
Optogenetic activation of a higher visual area and top-down effects in V1. **(A)** Illustration of the dorsal cortex of the cat indicating area boundaries (dashed lines). Virus was injected into the higher visual area 21a (green), and the effects of top-down activation were recorded in area 17 (purple). **(B)** Schematic of the visual cortical areas in the cat, depicting the major visual areas, their interconnections and relative hierarchical positions (top), as well as the laminar distribution of the dominant feed-forward and feed-back projections between areas 21a and 17. **(C)** Histology demonstrating the expression of EYFP. **(D)** Activation of Chr2 with blue light drives action potentials (blue trace in top panel) and gamma-band resonance in area 21a (blue trace in bottom panel). Control application of yellow light does not drive action potentials or gamma (green traces in the respective panels). **(E)** Optogenetic activation of area 21a combined with laminar recording in area 17 (both illustrated on the left), in the absence of visual stimulation. **(E, i)** Illustration of optogenetic stimulation and recording situation. **(E, ii)** Number of recording sites that showed significant OptoTD modulations of their firing rate in the absence of visual stimulation, as a function of cortical depth. Increases (decreases) are shown in green (red), and their sum in black. **(E,iii)** Firing rate changes induced by OptoTD in the absence of visual stimulation. Green (red) trace shows average over sites with significant increases (decreases), black trace the average over all significantly modulated sites, and blue trace the average over all sites. **(E, iv)** Magnitude of LFP power changes, averaged over all sites, irrespective of the sign of the change, as a function of frequency and cortical depth. Non-significant pixels are gray-shaded (FDR corrected for multiple comparisons).

To experimentally isolate the effects of area 21a on the primary visual cortex, we optogenetically stimulated^56–58^ excitatory neurons in area 21a, while recording populations of neurons in area 17. We report pronounced changes in gamma band synchronization that depended on features of the visual stimulus. Intriguingly, strong changes in synchronization could occur in the near absence of changes in firing rates of the recorded neurons and were not correlated to firing-rate changes. We additionally found that optogenetically controlled top-down modulation increased stimulus information encoded by populations of primary visual cortex neurons, and that these increases in stimulus information correlated specifically with changes in gamma-band synchronization, but not with changes in firing rate or synchronization in other frequency ranges.

## Results

### Effects of OptoTD on area 17 in the absence of visual stimulation

Viral transfection in area 21a was effective in all animals, as histology demonstrated the expression of the fluorophore coupled to the opsin (Fig. 1C for one example). Activation of the opsin through application of blue (473 nm) laser light induced strong and sustained increases in MUA and gamma, whereas the control application of yellow (594 nm) light did not (Fig. 1D).

We investigated the effect of optogenetic stimulation of area 21a (OptoTD) on neuronal activity in area 17, first in the absence of visual stimulation. OptoTD induced significant area 17 MUA modulation in less than 10% of sites (Fig. 1Ei, 3.5% increased firing rate (FR), 3.1% decreased FR). Significant increases occurred predominantly in deep layers, significant decreases in superficial layers (green and orange line, respectively, in Fig. 1Eii). Significant modulations overall showed a relative sparing of the granular layer (black line in Fig. 1Eii). When overall MUA modulation was expressed in the OptoTD-related difference in firing rate (black line in Fig. 1Eiii), differences ranged between approximately −2 and +1,5 spk/s.

For the further analysis, we quantified OptoTD-related modulations in MUA rate, LFP power or MUA-LFP pairwise phase consistency (PPC, see Methods), by calculating the OptoTD modulation index as

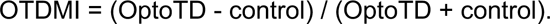

OptoTD induced significant area 17 LFP power modulation in several layers and frequencies (Fig. 1Eiv). In the lowest frequencies, the modulation extended over all layers. In the gamma band, the modulation occurred primarily in infragranular layers and most prominently in supragranular layers. Note that Fig. 1Eiv shows the OTDMI magnitude, irrespective of its sign, because positive and negative modulations were approximately balanced across sites (see below for a detailed analysis).

### Effects of OptoTD on area 17 in the presence of visual stimulation

The focus of this study was on the interaction of optogenetic top-down modulation with visual bottom-up drive. Visual stimuli were full-field and full-contrast moving gratings (lower contrast used only in Fig. 6) that induced layer-specific MUA and gamma-band synchronization as shown previously (Fig. S1).

Visually induced gamma-band synchronization often showed strong OptoTD modulation, either increases or decreases. Fig. 2A shows 16 example sites randomly selected from all sites with significant OptoTD-related increases in MUA-LFP gamma PPC, and Fig. 2B shows the same for gamma decreases. Note that for most of these sites, the OptoTD-related MUA modulation is very small, despite the large modulation of gamma synchronization.

**Figure 2:**
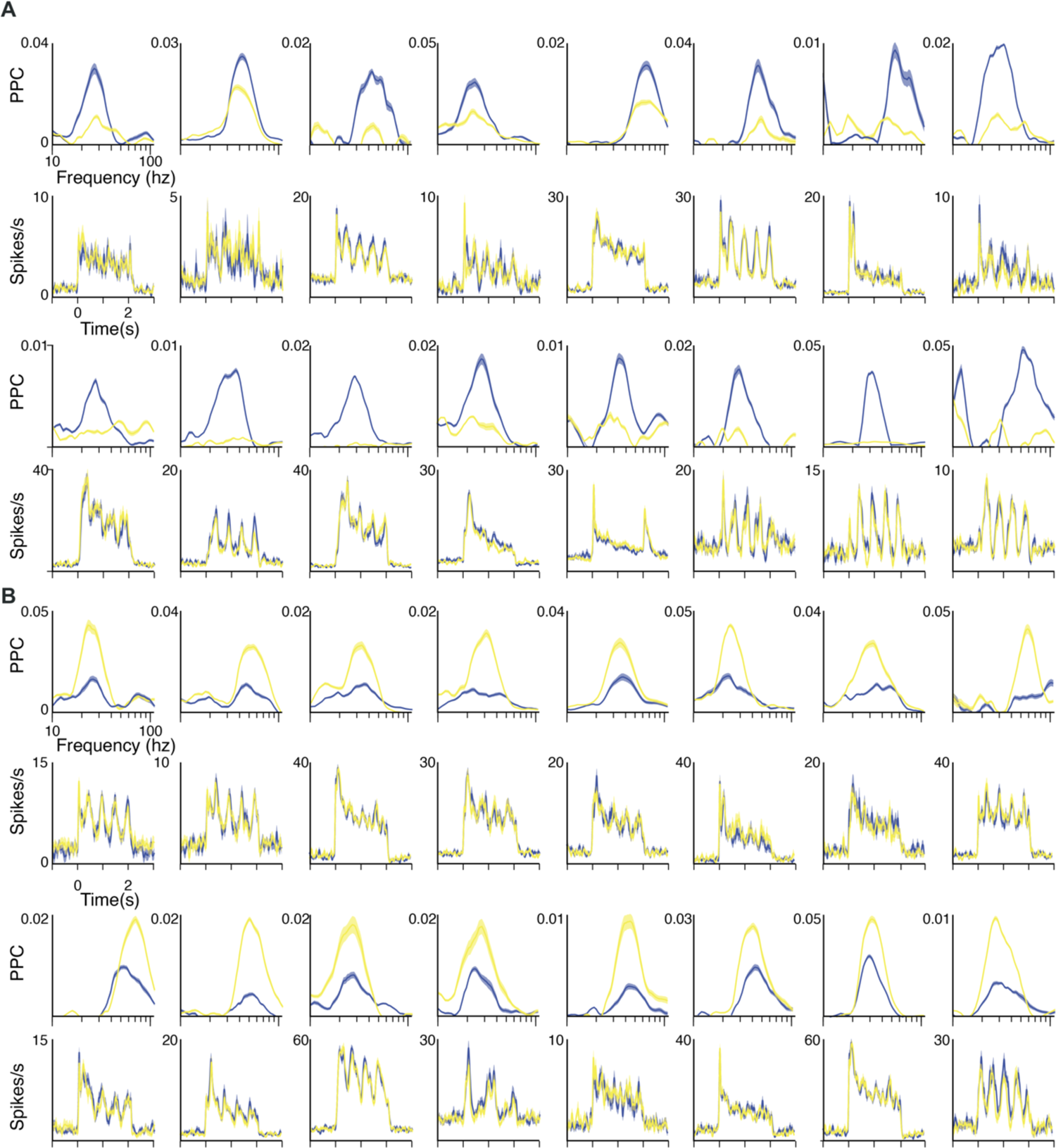
Top-down modulations of gamma-band synchronization in the primary visual cortex. **(A)** MUA-LFP PPC spectra and PSTHs for 16 example recording sites. For each site, the PPC spectrum is shown above the corresponding PSTH. Blue: OptoTD, yellow: Control. The 16 recording sites were randomly selected from all recording sites that showed significantly stronger gamma PPC for OptoTD versus Control. **(B)** Same as **(A)**, but for 16 recording sites that were randomly selected from all recording sites that showed significantly weaker gamma PPC for OptoTD versus Control. PPC spectra are clipped at zero as the lowest meaningful value.

To investigate this further, Fig. 3A shows 16 example sites randomly selected from all sites with significant OptoTD-related increases in MUA, and Fig. 3B shows the same for MUA decreases. Note that the corresponding modulations in MUA-LFP gamma PPC vary widely, with both increases and decreases.

**Figure 3.**
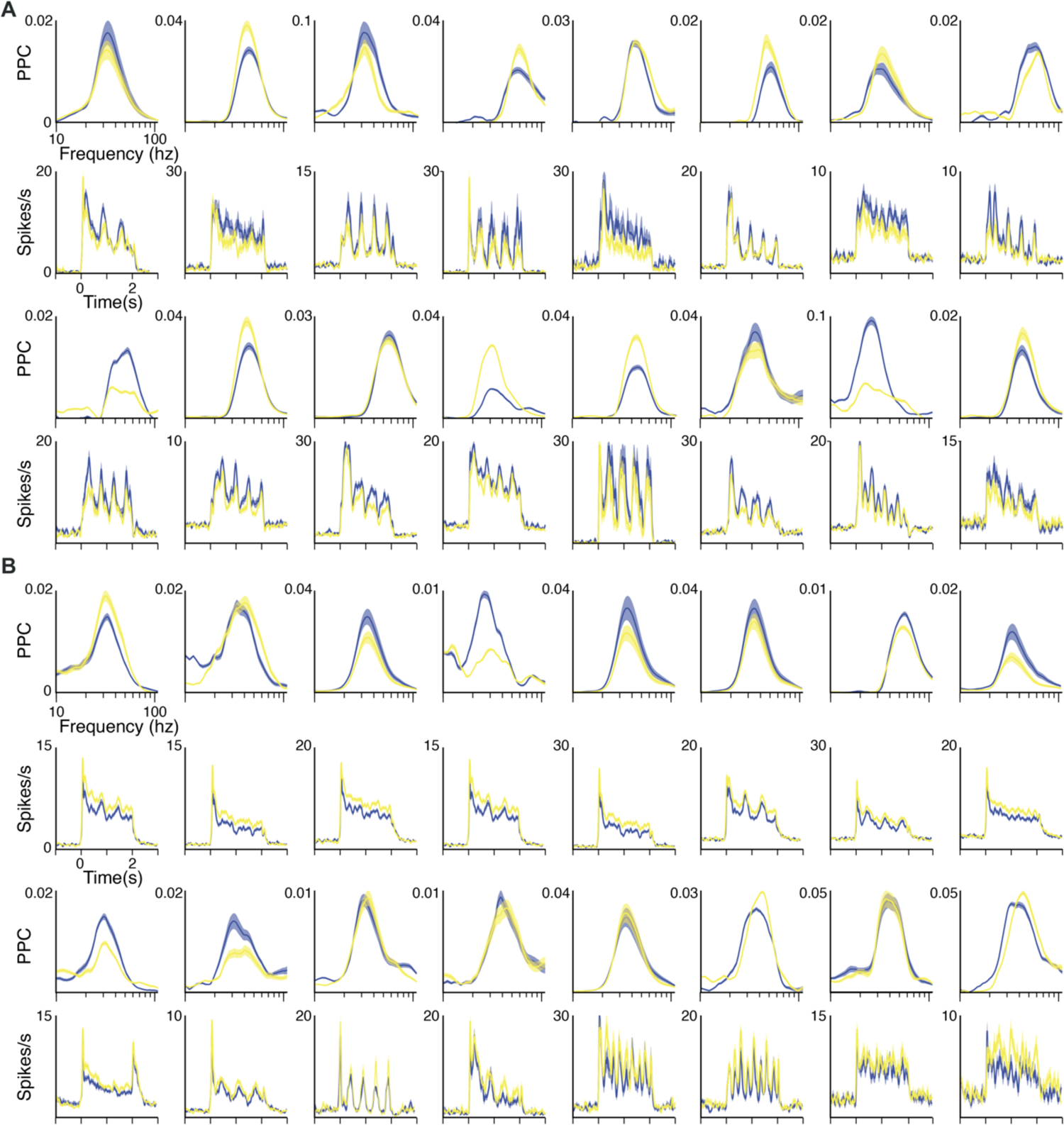
Top-down modulations of firing rates in the primary visual cortex. **(A)** MUA-LFP PPC spectra and PSTHs for 16 example recording sites. For each site, the PPC spectrum is shown above the corresponding PSTH. Blue: OptoTD, yellow: Control. The sites were randomly selected from all recording sites that showed significantly higher firing rates for OptoTD versus Control. **(B)** Same as **(A)**, but for 16 recording sites that were randomly selected from all sites that showed significantly lower firing rates for OptoTD versus Control. PPC spectra are clipped at zero, the lowest meaningful value.

We investigated these effects across all combinations of recording sites with visual stimulation conditions (i.e. stimulus directions), and separately for MUA, gamma- and low-frequency PPC. We found that the modulation magnitudes (increases or decreases) were much weaker for MUA rate than for low-frequency PPC or for gamma PPC (Fig. 4A,B; p<1^-100^ for both FR vs Low PPC and FR vs Gamma PPC, two-sample Kolmogorov-Smirnov test here and in the following; FR: mean=0.035, range=-0.43-0.42; Gamma: mean=0.23, range=-0.92-0.87; Low: mean=0.14, range=-0.74-0.67). The modulation was slightly stronger for gamma PPC than for low-frequency PPC (Fig. 4C; p<1^-80^). Fig. 4D superimposes the three OTDMI distributions for a direct comparison.

**Figure 4:**
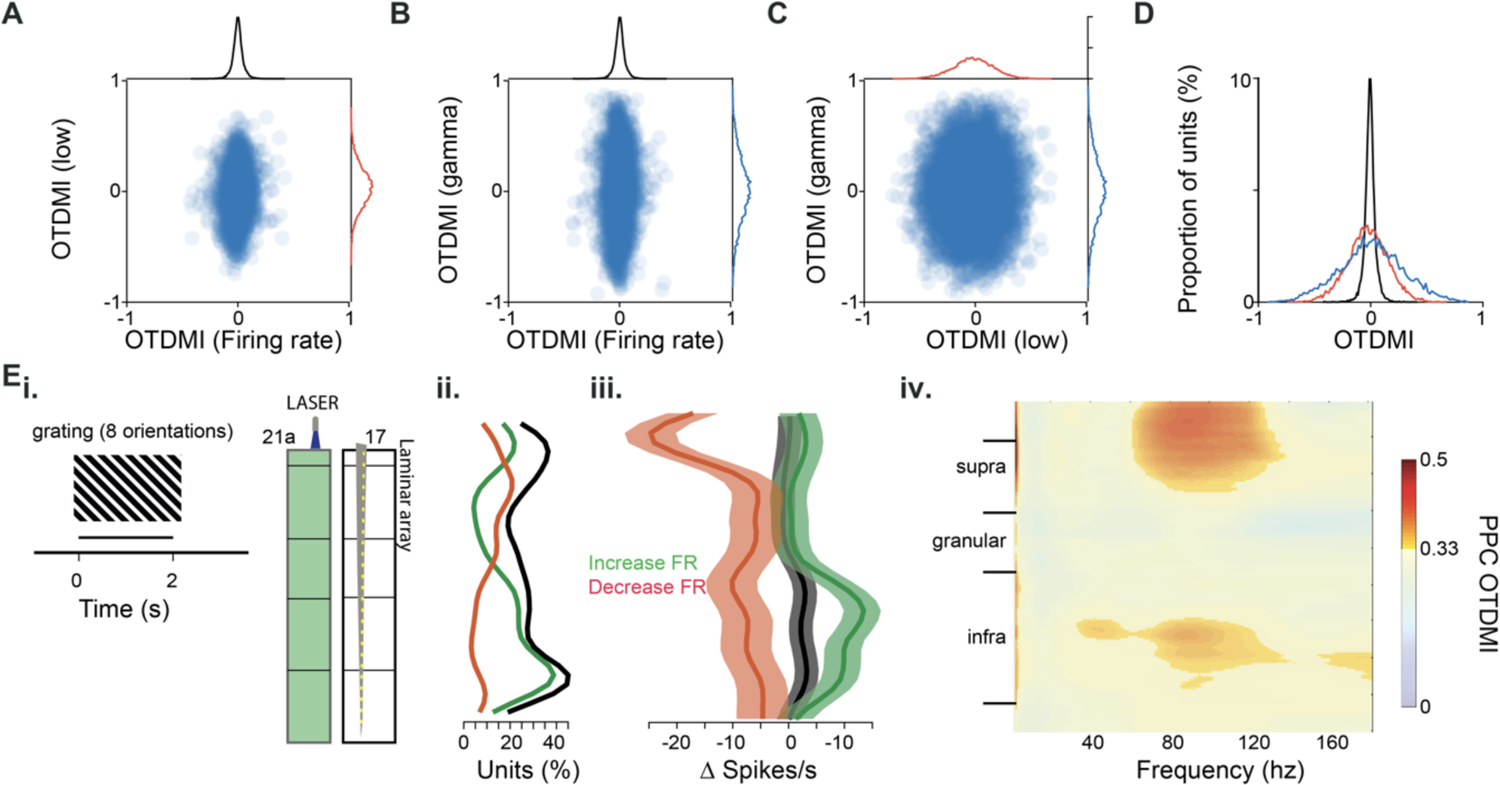
Effect of top-down modulation across the population. **(A)** Scatter plot of OptoTD-related modulations in MUA rate (x-axis) versus low-frequency MUA-LFP PPC (y-axis). Each dot corresponds to a combination of a recording site with a stimulus orientation. The x- and y-axis values are OptoTD indices defined as OI = (OptoTD – Control) / (OptoTD + Control). **(B)** Same as **(A)**, but for MUA rate (x-axis) versus gamma MUA-LFP PPC (y-axis). **(C)** Same as **(A)**, but for low-frequency MUA-LFP PPC (x-axis) versus gamma MUA-LFP PPC (y-axis). **(D)** Distributions of OptoTD indices. Same as the marginals plotted in (A-C), but superimposed to aid direct comparison. **(E)** Optogenetic activation of area 21a combined with laminar recording in area 17, in the presence of visual stimulation. **(E, i)** Illustration of optogenetic stimulation and recording situation. **(E, ii)** Number of recording sites that showed significant OptoTD modulations of their firing rate in the presence of visual stimulation, as a function of cortical depth. Increases (decreases) are shown in green (red), and their sum in black. **(E,iii)** Firing rate changes induced by OptoTD in the presence of visual stimulation. Green (red) trace shows average over sites with significant increases (decreases), black trace the average over all significantly modulated sites, and blue trace the average over all sites. **(E, iv)** Magnitude of LFP power changes, averaged over all sites, irrespective of the sign of the change, as a function of frequency and cortical depth. Non-significant pixels are gray-shaded (FDR corrected for multiple comparisons).

Interestingly, while there was a mild correlation between the OTDMI for low frequency PPC and firing rates, there was no correlation between the OTDMIs for gamma-band PPC and firing rates, suggesting that top-down signals can modulate synchronization in different frequency bands independently of firing rates (Spearman rank correlation: low PPC vs FR, rho=0.08 p<1^-15^; gamma PPC vs FR, rho= 0.01, p=0.2). OTDMIs for alpha-band PPC and gamma-band PPC showed a significant but very small correlation (Spearman rank correlation: low PPC vs gamma PPC, rho=0.02 p=0.025). A greater proportion of units was modulated by top-down activation when combined with visual stimulation (Fig. 4Ei-iii) than without (Fig. 1E; t-test on proportion, all: mean=26.6% vs 7.3%, p<1^-10^; increase: mean=16.3% vs 3.5%, p<1^-10^; decrease: mean=5% vs 3.1%, p<1^-10^), and OptoTD-induced firing rate changes were larger on average (Fig. 4Eiii, t-test on FR difference, all: mean=3.2 vs 1.9 spk/s, p<1^-20^, increase: mean=12.1 vs 9.6 spk/s, p<1^-12^, decrease: mean=-8.3 vs −6.1 spk/s, p<1^-13^). Despite the stronger OptoTD modulation when paired with visual stimulation, the laminar distribution of units with increased or decreased firing rates was highly similar (Fig. 4Eii-iii; Spearman-rank correlation across laminar distribution of proportion, all: rho=0.35, p=0.012; increase: rho=0.78, p<1^-8^; decreases: rho=0.77, p=1^-7^, Spearman-rank correlation across laminar distribution of FR difference, all: rho=0.38, p=0.02; increase: rho=0.70, p<1^-5^; decreases: rho=0.67, p=1^-6^). The OptoTD-induced PPC changes (Fig. 4Eiv) were likewise stronger with visual stimulation (mean=0.4 vs 0.31, t-test p1^-8^), but were highly similar in laminar and frequency distribution (Spearmann-rank correlation rho=0.42, p<1^-20^).

### OptoTD modulation depends on stimulus features

Given that OptoTD had both positive and negative effects on synchronization and firing rates, we next investigated whether these changes were systematically related to the visual stimulation. We found that the effects of OptoTD on synchronization and firing rate of individual sites depended on the features of the visual stimulus (Fig. 5). For some recording sites, OptoTD increased or decreased gamma-band synchronization in a direction-specific manner, increasing gamma-band synchronization for some stimulus directions, while decreasing it for others (Fig. 5A). For other recording sites, OptoTD increased (Fig. 5B), or decreased (Fig. 5C) gamma-band synchronization in an orientation-specific manner.

**Figure 5:**
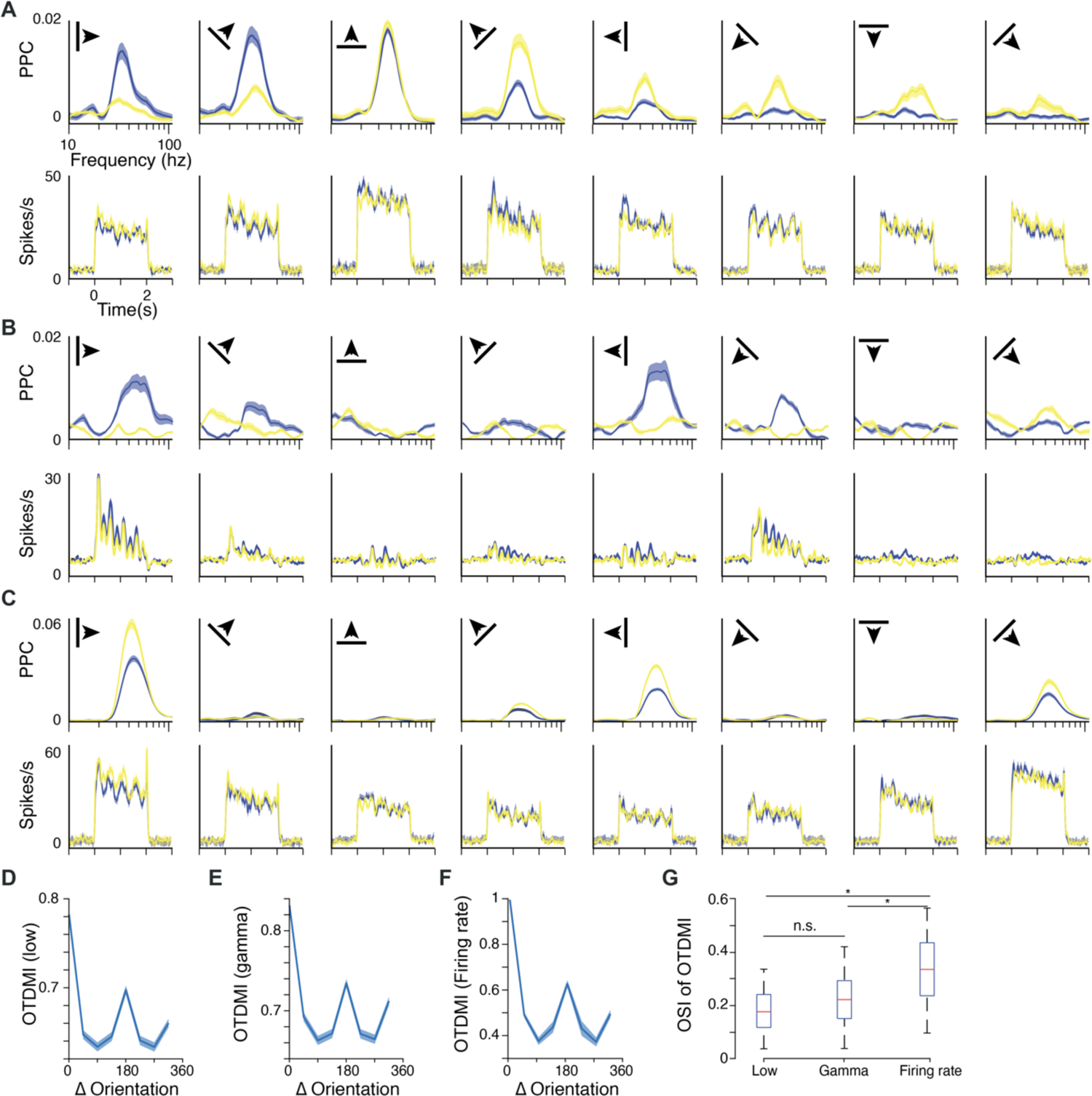
The effect of optogenetically induced cortical feedback depends on sensory stimulation. **(A)** MUA-LFP PPC spectra and PSTHs for one example recording site that showed OptoTD effects on the gamma-MUA PPC with a clear dependence on grating movement direction. For each stimulus orientation, the PPC spectrum is shown above the corresponding PSTH. Blue: OptoTD, yellow: Control. **(B)** Same as **(A)**, but for one example recording site that showed OptoTD-induced increases in the gamma-MUA PPC with a clear dependence on grating orientation. **(C)** Same as **(B)**, but for a recording site that showed OptoTD-induced decreases in the gamma-MUA PPC with a clear dependence on grating orientation. **(D)** Orientation tuning curves of OptoTD effect. **(D, i)** Rectified difference in low-frequency MUA-LFP PPC between OptoTD and Control, as a function of stimulus orientation. **(E)** Same as **(D)**, but for gamma MUA-LFP PPC. **(F)** Same as **(D)**, but for MUA rate. **(G)** Distribution of orientation selectivity index estimated on the OTDMI for low-frequency PPC, gamma PPC and MUA firing rates for each multiunit. PPC spectra are clipped at zero as the lowest meaningful value.

We explored the possibility that the OptoTD effect depended on the difference between the stimulus orientation and the preferred orientation of the recording site. Indeed, the OTDMI for the MUA-LFP PPC in the low-frequency range (Fig. 5D), the gamma range (Fig. 5E) and for the MUA rate (Fig. 5F) all showed clear tuning, with a main peak at the preferred orientation and a secondary peak 180 deg from the preferred orientation. Indeed, estimating the orientation selectivity index (OSI) of the OTDMI for synchrony and firing rate for each multiunit indicated that the influence of top-down signals on primary visual cortex neurons had significant stimulus tuning (Fig. 5G) that was similar, but significantly stronger, for MUA FR than for low-frequency and gamma synchrony (t-test, p<0.03 for OSI of OTDMI for MUA FR vs both low-frequency and gamma PPC).

### OptoTD increases stimulus information in V1

The stimulus-specificity of OptoTD effects led us to ask whether OptoTD influenced stimulus encoding in the V1 population. To this end, we trained classifiers to decode the direction of motion of the visual stimulus either from single MUA recording sites or from groups of simultaneously recorded MUA sites (labeled “population” in Fig. 6). We found that OptoTD could increase decoder accuracy (Fig. 6A presents population MUA decoding from a single example session), and over all multi-units across sessions increased single-MUA accuracy by 3.6% on average (range: −5.3% – 14.1%), and population accuracy by 6.7% on average (range: 1.2%-14.3%) (Fig. 6B). OptoTD-related increases in decoder accuracy were larger in the deep layers (Fig. 6C, t-test, p<0.012) and increased as stimulus contrast decreased (Fig. 6D, t-test, * denotes p<0.02 and ** denotes p<0.01), suggesting that OptoTD could increase the stimulus information for weak stimulus drive. OptoTD-related increases in accuracy correlated with OptoTD-related changes specifically in gamma-band synchronization, but not in low-frequency synchronization or MUA rate (Fig. 6 E, Spearman-rank correlation between accuracy and: OTDMI FR, rho=0.32, p=0.061; low frequency PPC, rho=-0.01, p=0.92, gamma frequency PPC, rho=0.68, p<1^-6^).

**Figure 6:**
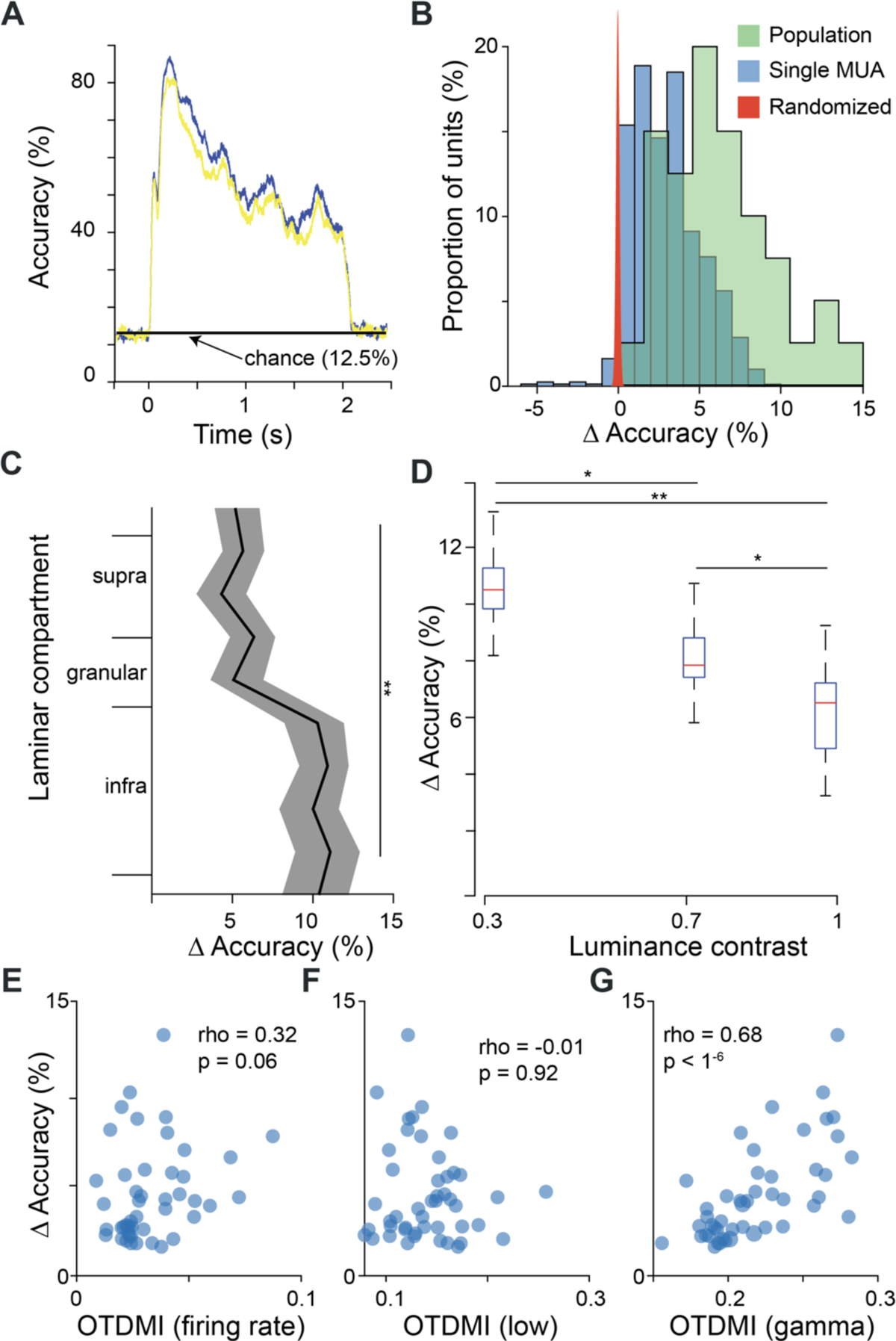
Top-down modulation improves MUA-rate-based stimulus decoding in primary visual cortex. **(A)** Accuracy of MUA-rate-based stimulus decoding as a function of time relative to stimulus onset, decoder based on population vector across MUAs for one example session. Blue: OptoTD, yellow: Control. **(B)** Blue: Distribution of differences in accuracy between OptoTD and Control (averaged over the entire visual stimulation period) for single MUA decoding across all multiunits and sessions. Green: Same distribution, but for population decoding across all sessions. Red: Population decoding distribution after randomly shuffling OptoTD versus Control. **(C)** Difference (not rectified) in accuracy between OptoTD and Control, averaged over the entire visual stimulation period and over all session ensembles, as a function of cortical depth. **(D)** Difference (not rectified) in accuracy between OptoTD and Control, averaged over the entire visual stimulation period and over all session ensembles, as a function of visual stimulus contrast. Note that all other analyses use full contrast. (* denotes p<0.02 and ** denotes p<0.01) **(E)** Scatter plot of OptoTD-related differences in MUA-rate-based decoding accuracy versus MUA rate. **(F)** Same as **(E)**, but versus low-frequency MUA-LFP PPC. **(G)** Same as **(E)**, but versus gamma MUA-LFP PPC.

## Discussion

Neurons in early sensory areas do not simply represent the world in a fixed manner but must integrate current sensory information with intrinsic activity arising from local and long-range connections. One prominent source of long-range connections to the primary visual cortex are the feedback projections that transmit information down the visual hierarchy from higher-order visual areas in a top-down manner. In this study, we used optogenetics to control top-down signals from area 21a, a higher-order area in the cat ventral visual stream, while recording the activity of populations of neurons in the primary visual cortex. We found that optogenetic activation of area 21a during simultaneous visual stimulation had the largest effect on gamma-band synchronization, with relatively minor, and independent effects on the firing rates of neurons in area 17. These modulations exhibited a distinct laminar profile that matches the anatomy of feedback connections in the visual hierarchy. Surprisingly, changes in firing rates and synchronization were uncorrelated, and large top-down mediated increases or decreases of gamma-band synchronization could occur in the absence of firing rate changes. Both gamma-band and firing-rate changes depended on the features of the visual stimulus, preserving, and enhancing the stimulus selectivity of area 17 rather than overwriting it. OptoTD also increased the fidelity of firing-rate responses and the accuracy of decoders trained to predict stimulus identity. The magnitude of these increases was dependent on the visual contrast of the sensory input, with larger OptoTD effects for weaker stimuli. The magnitude of increased decoding performance correlated with OptoTD-induced changes in gamma-band synchronization, but not with changes in firing rates, or low-frequency synchronization. Overall, our results indicate that top-down signals within the visual hierarchy may dominantly modulate the gamma-band synchronization of early sensory neurons, while sparing other coding channels, such as firing rates or synchronization in other frequency ranges to accurately represent the world or convey independent sources of top-down information.

While the results of our study are clear, our experiments had several limitations that may constrain the specificity or generality of our findings. First, the expression of ChR2 in area 21a was performed with broad viral injections and was agnostic to any underlying functional or anatomical organization. While we limited expression to excitatory neurons using the CamKII promoter, we did not target a specific functional or anatomical subpopulation, for example, according to stimulus preference or projection target. As such, our OptoTD protocol most likely engaged many diverse excitatory neurons and so may have activated area 21a in a less than fully physiological way. Future studies could improve upon this by limiting expression to a specific functional subclass of higher-order neurons, to only 21a neurons that project to area 17, or further by targeted activation of functional ensembles using single-cell perturbation methods. Second, the visual stimulation used in our study was limited to full-field gratings, visual stimuli which strongly activate neurons in the primary visual cortex and are known to produce prominent gamma-band synchronization, but which are unnatural, and which lead to strong lateral interactions within area 17. While this stimulus set is unnatural, it is commonly used in visual experiments because it provides parametric control of the input and strongly activates the neurons in the primary visual cortex. Future studies may use more refined stimulus sets to further examine the potentially rich coding capacity of top-down projections within the visual hierarchy, and to investigate the effects of top-down signals on lateral integration in area 17. Finally, our experiments were performed in anesthetized cats rather than awake, behaving subjects. While many anesthesia protocols have prominent effects on neurons and sensory responses, we used a hybrid anesthesia protocol based on a potent opioid supplemented by very low doses of isoflurane, resulting in a light anesthesia with visual responses and neuronal rhythms comparable to the awake animal. Further, we believe that anesthesia may be a feature of the present study, as it allowed us to suppress many ongoing processes, which may have engaged diverse sources of top-down signals to the early visual cortex. In this case, anesthesia may have enabled us to cleanly activate our top-down source of interest, in the absence of competing and confounding top-down signals. That being said, future studies should further investigate the compelling observations we have made in the anesthetized preparation and see how top-down modulation of gamma-band synchronization affects sensory encoding, information transmission and perceptual performance in awake, behaving animals.

Other studies have used direct manipulation of specific brain regions to investigate top-down processes in the awake or anesthetized brain^44,45,59–65.^ These studies have complemented anatomical studies to demonstrate that long-range cortical projections can have functional effects in early visual areas, focusing mostly on the effect of top-down cortico-cortical signals on firing rates and perception. It has also been shown that suppression of top-down signals (cooling area MT in the primate) can reduce visual responses and figure-ground segmentation in V1^66^. Two recent studies have investigated the effects of suppressing top-down signals by cortical cooling or optogenetic inhibition of intermediate or higher order visual areas (V2 or V4) in the primate visual hierarchy while assessing the effects on synchronization in V1^59,60^. Both studies reported decreases in gamma-band synchronization when suppressing top-down signals in combination with visual stimulation. These results are broadly consistent with our results, though we found that optogenetic activation of the higher areas could both increase or decrease gamma-band synchronization for different recording sites and dependent on the simultaneously applied visual stimulus.

While our study was unable to reveal the precise cellular or network mechanisms giving rise to our observed OptoTD effects, recent work in the mouse may shed some light on the underlying mechanisms. The development of powerful genetic and experimental approaches available in the mouse have enabled the exploration of the circuit mechanisms of top-down signals arising from specific cortical feedback projections. For example, the direct projection from the Anterior cingulate cortex to the primary visual cortex of mice (a connection that does not exist in primates) has been shown to enhance orientation tuning and center-surround interactions through projections onto local inhibitory interneurons in V1^67^. Using an all-optical, single-cell perturbation approach, Fişek and colleagues have recently demonstrated the effect of top-down signals from higher-order visual cortex onto neurons in the primary visual cortex of mice that can enhance or suppress the activity of V1 neurons depending on the precise retinotopic alignment of the neurons^68^.

Critical to our findings are the precise local and long-range circuits that can enable OptoTD to powerfully modulate gamma-band synchronization independently of the firing rate of neurons. One possibility is suggested by work in mice that demonstrates long-range connections specifically targeting vasoactive intestinal polypeptid (VIP) positive interneurons in the primary visual cortex^67^. VIP interneurons are known to demonstrate strong modulation by arousal and behavioral state^69^, and their activity correlates strongly with positive and negative reinforcement signals^70^, believed to arise from long-range, top-down projections and subcortical structures. Interestingly, VIP interneurons appear to synapse preferentially onto somatostatin-expressing interneurons (SOM), which in turn inhibit both parvalbumin positive interneurons (PV) and pyramidal neurons (PYR)^71^. Intriguingly, the tight interplay between PYR and neighboring PV neurons characteristic of cortical circuits has been well-documented to be essential to the generation of cortical gamma-band synchronization^72–76^. A prominent gamma-band resonance has been extensively documented in the cat visual system^77–80^, and we recently used optogenetics to probe the effects of this resonance on signal transmission in the visual hierarchy^57,58^. In such a scenario, top-down signals from the visual hierarchy might simultaneously modulate PYR and PV neurons and thereby gamma-band synchronization in V1 without concomitantly altering the firing rate or stimulus tuning of V1 neurons.

Indeed, this scenario might provide intriguing functional opportunities: Top-down signals might dynamically alter the plasticity, coordination, and routing of information in a flexible way depending on goals and context, without affecting the accurate encoding of low-level sensory information^81–83^. In this framework, the firing rates of V1 neurons would primarily provide a veridical reflection of the external world, minimally corrupted by moment-to-moment variations in the animal’s drives, goals, or behavioral states^84,85^. At the same time, gamma-band synchronization would primarily serve the selective routing of V1 information based on behavioral control of attention^2,51,52,86,87^, or promote plasticity of behaviorally relevant stimuli ^88–91^. We would like to speculate that this might be an evolutionary development that has emerged specifically in higher mammals. Humans, non-human primates and cats show particularly strong gamma-band synchronization in primary visual cortex. At the same time, effects of behavioral and cognitive state on firing rates in the visual system typically decrease with decreasing hierarchical level of the investigated area and are smallest in primate area V1^92–94^. While an inter-species comparison is difficult due to differences in tasks and other factors, effect sizes found in mid- to high-level primate visual areas might be most comparable to those found in rodent V1, where firing rates can be strongly affected by top-down signals such as arousal and motor activity^20,95,96^. Consistent with this, V1 receives stronger and longer-range direct top-down projections in rodents than in primates^32,97^, and primate visual-area anatomical connectivity reveals more segregation and modularization^36,98^. Such a separation could be specific to strongly visual animals, such as cats and primates, for which the visual system is more hierarchically structured.

It will be interesting for future studies to further refine the investigation of effects that top-down signals across the visual hierarchy can have on processing in primary visual cortex. One notable feature of top-down signals in the awake animal is their temporal structure^65,99–105^, a dimension which we did not probe in our study, but which future studies should investigate. Likewise, experiments that limit opsin expression to neurons with specific anatomical, genetic, or functional features will continue to reveal the intricate patterns of information conveyed down sensory hierarchies.

## Acknowledgements

We thank Jianguang Ni for assistance during a portion of the experiments reported here. This work was supported by DFG (SPP 1665 FR2557/1-1, FOR 1847 FR2557/2-1, FR2557/5-1-CORNET, FR2557/6-1-NeuroTMR, FR2557/7-1-DualStreams to P.F.), FP7-604102-HBP, FP7-600730-Magnetrodes to P.F., National Institutes of Health (1U54MH091657-WU-Minn-Consortium-HCP to P.F.), the LOEWE program (NeFF to P.F.).

## Author contributions

C.M.L., T.W., and P.F. designed research; C.M.L. and T.W. performed experiments; C.M.L. analyzed data and prepared figures in consultation with P.F.; C.M.L. and P.F. wrote the paper; P.F. supervised the project.

## Methods

### RESOURCE AVAILABILITY

#### Lead contact

Further information and requests for resources and reagents should be directed to and will be fulfilled by the Lead Contact, Christopher Lewis.

#### Materials availability

This study did not generate new unique reagents.

#### Data and code availability

The datasets and code supporting the current study are available from the corresponding author on request.

#### EXPERIMENTAL MODEL AND SUBJECT DETAILS

A total of 67 sessions from four adult domestic cats (*felis catus*; two females; mean age 4.2 year; range 3-8 years, 15-20 sessions per cat) were used in this study. We used cats because the visual hierarchy is well developed, and the visual cortical physiology is highly similar to human and non-human primates, both during wakefulness and light anesthesia. Different data from the same animals were used in previous studies^57,58,106^. All procedures complied with the German law for the protection of animals and were approved by the regional authority (Regierungspräsidium Darmstadt). After an initial surgery for the injection of viral vectors and a 4-6 week period for opsin expression, recordings were obtained during a terminal experiment under general anesthesia.

## METHOD DETAILS

### Viral vector injection

For the injection surgery, anesthesia was induced by intramuscular injection of ketamine (10 mg/kg) and dexmedetomidine (0.02 mg/kg), cats were intubated, and anesthesia was maintained with N_2_O:O_2_ (60/40%), isoflurane (∼1.5%) and remifentanil (0.3 µg/kg/min). All four cats were injected in area 21a. Rectangular craniotomies were made over the area of injection to identify area 21a by local anatomical landmarks (AP [0,-8] mm, ML [9, 15] mm). The injection sites were chosen based on the pattern of sulci and gyri, and the dura mater was removed over part of the respective areas. Three to four injection sites were chosen, avoiding blood vessels, with horizontal distances between injection sites of at least 1 mm. At each site, a Hamilton syringe (34 G needle size; World Precision Instruments) was inserted with the use of a micromanipulator and under visual inspection to a cortical depth of 1 mm below the pia mater. Subsequently, 2 µl of viral vector dispersion was injected at a rate of 150 nl/min. After each injection, the needle was left in place for 10 min before withdrawal, to avoid reflux. Upon completion of injections, the dura opening was covered with silicone foil and a thin layer of silicone gel, the trepanation was filled with dental acrylic, and the scalp was sutured.

In each cat, area 21a of the left hemisphere was injected with AAV9-CamKIIα-hChR2(H134R)-eYFP^107^ (titer: 1.06*10^13^ GC/ml). The viral vector was obtained from Penn Vector Core (Perelman School of Medicine, University of Pennsylvania, USA).

### Neurophysiological recordings

For the recording experiment, anesthesia was induced and initially maintained as during the injection surgery, only replacing intubation with tracheotomy and remifentanyl with sufentanil. After surgical procedures were complete and prior to recordings, isoflurane concentration was lowered to 0.6%-1.0%, eye lid closure reflex was tested to verify narcosis, and vecuronium (0.25mg/kg/h i.v.) was added for paralysis during recordings. Throughout surgery and recordings, Ringer solution plus 10% glucose was given (20 ml/h during surgery; 7 ml/h during recordings), and vital parameters were monitored (ECG, body temperature, expiratory gases).

Each recording experiment consisted of multiple sessions. For each session, we inserted either multiple tungsten microelectrodes (∼1 MΩ at 1 kHz; FHC), or two to four 32-contact probes (100 µm inter-site spacing, ∼0.3 MΩ at 1 kHz; NeuroNexus or ATLAS Neuroengineering) in area 17. Standard electrophysiological techniques (Tucker Davis Technologies, TDT) were used to obtain multi-unit activity (MUA) and LFP recordings. For MUA recordings, the signals were filtered with a passband of 700 to 7000 Hz, and a threshold was set to retain the spike times of small clusters of units. For LFP recordings, the signals were filtered with a passband of 0.7 to 250 Hz and digitized at 1017.1 Hz.

### Photo-stimulation

We used a laser system with two wavelengths for optogenetic stimulation (473 nm, blue laser) or control (594 nm, yellow laser). Laser light was delivered to cortex through a 100 µm or a 200 µm diameter multimode fiber (Thorlabs). Fiber endings were placed just above the cortical surface in area 21a. The intensity and waveform of the laser were controlled via a custom circuit in TDT, and timing was controlled by the visual stimulus computer using Psychtoolbox-3, a toolbox in MATLAB (MathWorks)^108^.

### Histology

After conclusion of recordings, approximately five days after the start of the terminal experiment and still under narcosis, the animal was euthanized with pentobarbital sodium and transcardially perfused with phosphate buffered saline (PBS) followed by 4% paraformaldehyde. The brain was removed, post-fixed in 4% paraformaldehyde and subsequently soaked in 10%, 20% and 30% sucrose-PBS solution, respectively, until the tissue sank. The cortex was sectioned in 50 µm thick slices, which were mounted on glass slides, coverslipped with an antifade mounting medium, and subsequently investigated with a confocal laser scanning microscope (CLSM, Nikon C2 90i, Nikon Instruments) for eYFP-labelled neurons. We previously documented the specific transfection of layer 2/3 and layer 5/6 pyramidal neurons and the absence of inhibitory interneurons^57^. Critical to this study, we found no transfected neurons in area 17.

### Data analysis

Data analysis used the Fieldtrip toolbox^109^, written in MATLAB (MathWorks).

#### Current source density (CSD) estimation, LFP power spectra, and MUA-LFP synchrony

Estimation of the CSD from the multi-electrode local field potential recordings used a modified version of the inverse current source density method (Pettersen 2006). Calculation of LFP power spectra was based on data epochs that were adjusted for each frequency to have a length of 5 cycles and moved over the data in a sliding-window fashion in 10 ms steps. Each epoch was multiplied with a Hann taper, Fourier transformed, squared and divided by the window length to obtain power density per frequency. The LFP power ratio was calculated by dividing the power during stimulation (0.2 s to 0.5 s relative to stimulation onset) with the pre-stimulation baseline power (−0.5 s to −0.2 s relative to stimulation onset).

MUA-LFP locking was quantified by calculating the MUA-LFP PPC (pairwise phase consistency), a metric that is not biased by trial number, spike count or spike rate^110^. Spike and LFP recordings for MUA-LFP PPC analysis were always taken from different electrodes. For each spike, the surrounding LFP was Hann tapered and Fourier transformed. Per spike and frequency, this gave the MUA-LFP phase, which should be similar across spikes, if they are locked to the LFP. This phase similarity is quantified by the PPC as the average phase difference across all possible pairs of spikes. For a given MUA channel, MUA-LFP PPC was calculated relative to all LFPs from different electrodes and then averaged.

#### Spike rate

All MUA analysis was performed on the spike density calculated from all threshold-crossings on a given electrode. Spike density was computed on a single-trial basis by convolving the binary spike time-series with a Gaussian kernel (standard deviation of 10 ms). Each binary time-series was separately convolved with the kernel in order to derive a continuous variable, which divided by the time window, led to a time-varying spike density estimate.

#### Naïve-Bayes decoding

The spike density was estimated for each multiunit for each trial in 100 ms windows for each condition. For MUA decoding, either the responses from single MUA recording sites or the population vector from MUA recorded simultaneously across the multi-contact array was used to train a probabilistic decoder using the Gaussian naïve Bayes algorithm. Naïve Bayes applies Bayes’ Theorem with the assumption that features are independent to estimate P(X|Y), the probability of feature X, given condition Y. The feature vector was composed of a single time window across recording sites, for population analysis, or of a single MUA across time for single-MUA performance. The decoder estimates the mean and standard deviation of a Gaussian for each feature (MUA) and each condition (orientation or direction). When the trained distributions are applied to previously unseen data, the decoder calculates the likelihood associated with each of the possible feature classes (8 orientations, 16 directions of motion). The stimulus class is predicted to be the maximum likelihood across classes and compared to the veridical class to quantify classification accuracy. To compare decoding under different LASER conditions, we either pooled half of the training trials from the blue and yellow laser conditions, or we trained only on the yellow LASER condition and tested it on both LASER conditions. Because there was no qualitative difference in the results, we only present the results obtained by training on the pooled blue and yellow LASER conditions. The training set consisted of 50% of the available trials (half from the blue condition and half from the yellow), and performance was quantified on the remaining 50% of trials separately per LASER condition. We used an iterated cross-validation approach, which allowed us to control the generalization and robustness of our decoder and provide confidence intervals for decoder accuracy. Values displayed for the decoding are the mean value from 100 iterations of the decoder to avoid any extraneous effects of the training and test sets.

## Supporting Information

**Supplementary Figure 1:**
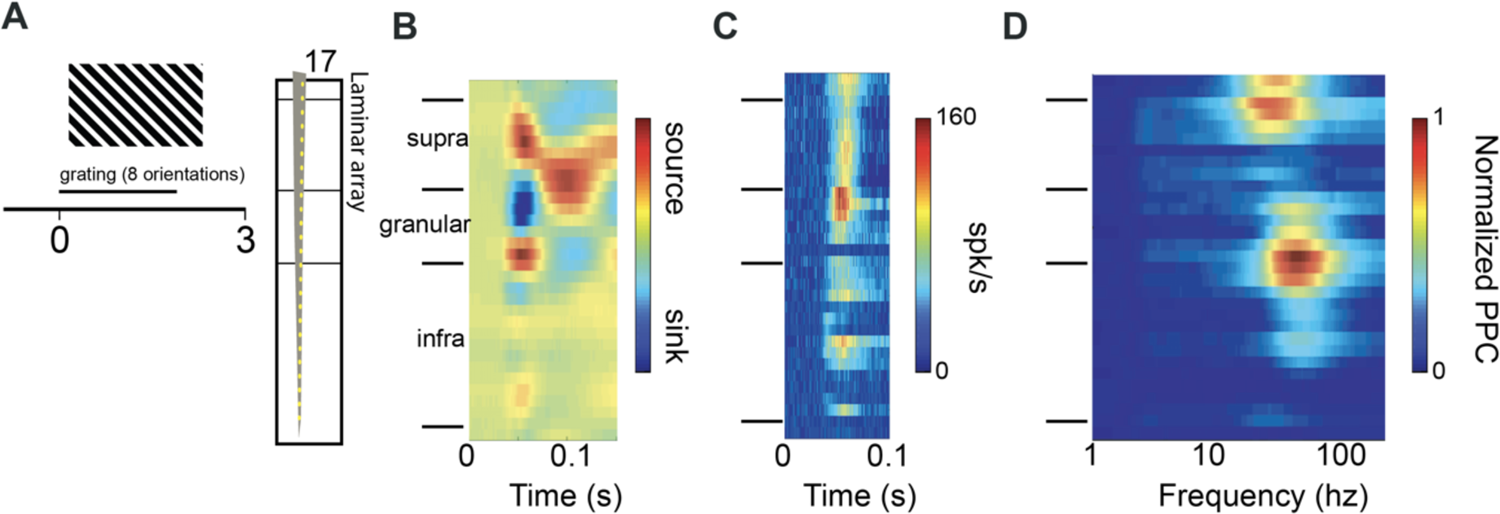
Visual stimulation induces layer-specific MUA and gamma-band synchronization. (A) Illustration of visual stimulation and laminar recording in area 17, in the absence of any optogenetic activation. Visual stimulation with a large, full-contrast grating (8 different orientations across trials). (B) Current-source density as a function of time after stimulus onset and cortical depth. (C) Firing rates as a function of time after stimulus onset and cortical depth. (D) MUA-LFP PPC as a function of frequency and cortical depth.

